# Sparsification of Large Ultrametric Matrices: Insights into the Microbial Tree of Life ^*^

**DOI:** 10.1101/2022.08.21.504697

**Authors:** Evan D. Gorman, Manuel E. Lladser

**Affiliations:** Department of Applied Mathematics, University of Colorado, Boulder, CO 80309, The United States

**Keywords:** Double Principal Coordinate Analysis, Haar-like wavelets, sparsification, phylogenetic covariance matrix, strictly ultrametric matrix, Tree of Life, UniFrac

## Abstract

Strictly ultrametric matrices appear in many domains of mathematics and science; nevertheless, they can be large and dense, making them difficult to store and manipulate, unlike large but sparse matrices. In this manuscript, we exploit that strictly ultrametric matrices can be represented as binary trees to sparsify them via an orthonormal base change based on Haar-like wavelets. We show that, with overwhelmingly high probability, only an asymptotically negligible fraction of the off-diagonal entries in random but large strictly ultrametric matrices remain non-zero after the base change; and develop an algorithm to sparsify such matrices directly from their tree representation. We also identify the subclass of matrices diagonalized by the Haar-like wavelets and supply a sufficient condition to approximate the spectrum of strictly ultrametric matrices outside this subclass. Our methods give computational access to the covariance matrix of the microbiologists’ Tree of Life, which was previously inaccessible due to its size, and motivate introducing a new wavelet-based (beta-diversity) metric to compare microbial environments. Unlike the established (beta-diversity) metrics, the new metric may be used to identify internal nodes (i.e., splits) in the Tree that link microbial composition and environmental factors in a statistically significant manner.

**MSC codes:** 05C05, 15A18, 42C40, 65F55, 92C70

## 1. Introduction

Ultrametric matrices appear across many domains of mathematics and science. They comprise an important class of matrices called inverse-M matrices [11] and are a key object of study in potential theory and Markov Chains [12]. In scientific applications, ultrametric matrices act as covariance models in phylogenetic comparative analysis [47], network inference [30], and energy models in statistical physics [8]. Further hinting at the pervasiveness of ultrametric matrices in modern data science, recent work has shown that the matrix of normalized Euclidean distances between points in some random subsets of ℝ^*d*^ converge in probability to an ultrametric matrix as *d* tends to infinity [61, 62].

In many applications, the underlying ultrametric matrix can be dense and potentially too large to store in computer memory and manipulate. Nonetheless, if a sparse representation of such a matrix can be found, many otherwise impossible tasks become computationally feasible, such as matrix inversion, eigenvalue decompositions, and principal component analysis (PCA).

This paper addresses the challenge of sparsifying the subclass of so-called strictly ultrametric matrices, which we define next.

In what remains of this manuscript, *n* ≥ 1 is an integer and [*n*] := {1, …, *n*}. Vectors and sometimes functions are represented as column vectors, and the transpose of a vector or matrix *A* is denoted *A*′.

### Definition 1.1 ([57]).

*A matrix S* ∈ ℝ^*n*×*n*^ *is ultrametric if it is symmetric with nonnegative entries and S*(*i, j*) ≥ min {*S*(*i, k*), *S*(*k, j*)} *for all i, j, k* ∈ [*n*]. *If in addition S*(*i, i*) *>* max {*S*(*i, t*) : *t* ≠ *i*}, *for all i* ∈ [*n*], *S is called strictly ultrametric. For n* = 1, *the last inequality is replaced with S*(*i, i*) *>* 0.

Ultrametric matrices have rich properties that are not made evident by their definition [11]. In particular, if *S* is strictly ultrametric then it is positive definite (hence invertible), *S*^−1^ is strictly diagonally dominant with non-positive off-diagonal entries, and *S*(*i, j*) = 0 if and only if *S*^−1^(*i, j*) = 0. These properties were initially proved using probabilistic methods [40]. An alternative proof is based on an equivalence between strictly ultrametric matrices and a subclass of binary trees [44]. The key ingredient for this equivalence is that for *n >* 1, if *S* ∈ ℝ^*n*×*n*^ is symmetric with non-negative entries then it is strictly ultrametric if and only if there exists a permutation matrix *P* and strictly ultrametric matrices *A* and *B* such that

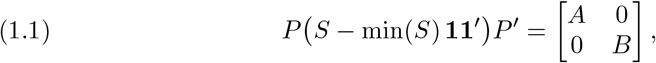

where min(*S*) is the smallest entry in *S*, and **1** ∈ ℝ^*n*^ is the column vector of ones [44, Proposition 2.1]. Since *A* and *B* are of the same kind as *S*, this process may be applied recursively and the matrix *S* encoded as a weighted rooted binary tree with special characteristics. Here we adopt a slightly different encoding to the one in [44], which is more suitable for our purposes. The reader unfamiliar with the jargon and notation of trees may skip ahead to Section 1.2 and come back to make better sense of the construction below.

Let *S* be a strictly ultrametric matrix of dimensions *n* × *n*. We can represent *S* as a rooted binary tree with 2*n* nodes (of which half are leaves) and hence (2*n* − 1) edges, satisfying the following definition.

### Definition 1.2.

*An out-rooted bifurcating tree (ORB-tree) with n leaves is a weighted rooted tree with the following properties: each vertex has degree* 1 *or* 3; *its leaf set is* [*n*] *and excludes the root, which has degree 1; each edge is labeled by the subset of leaves that descend from it; and the length ℓ*(*e*) *of each edge e is non-negative but ℓ*(*e*) *>* 0 *when e connects a leaf with its parent*.

The representation of a strictly ultrametric matrix *S* as an ORB-tree may be obtained as follows. The only edge emanating from the root is labeled as [*n*] and defined to have length min(*S*). The only child of the root has two children. One child descends from an edge labeled by the rows (or columns) of *S* associated with the matrix *A* before applying the permutation matrix *P* in (1.1). This edge has length min(*A*). Likewise, the other child descends from an edge labeled by the rows associated with the matrix *B* and has length min(*B*). Since *A* and *B* are strictly ultrametric, just of smaller dimensions, the tree may be grown recursively from any descendent of the root that is not associated with a strictly ultrametric matrix of dimensions 1 × 1. The latter represent edges that parent a leaf in the ORB-tree. These edges must have a strictly positive length because 1 × 1 strictly ultrametric matrices are strictly positive real numbers. To fix ideas see Figure 1.

**Fig. 1:**
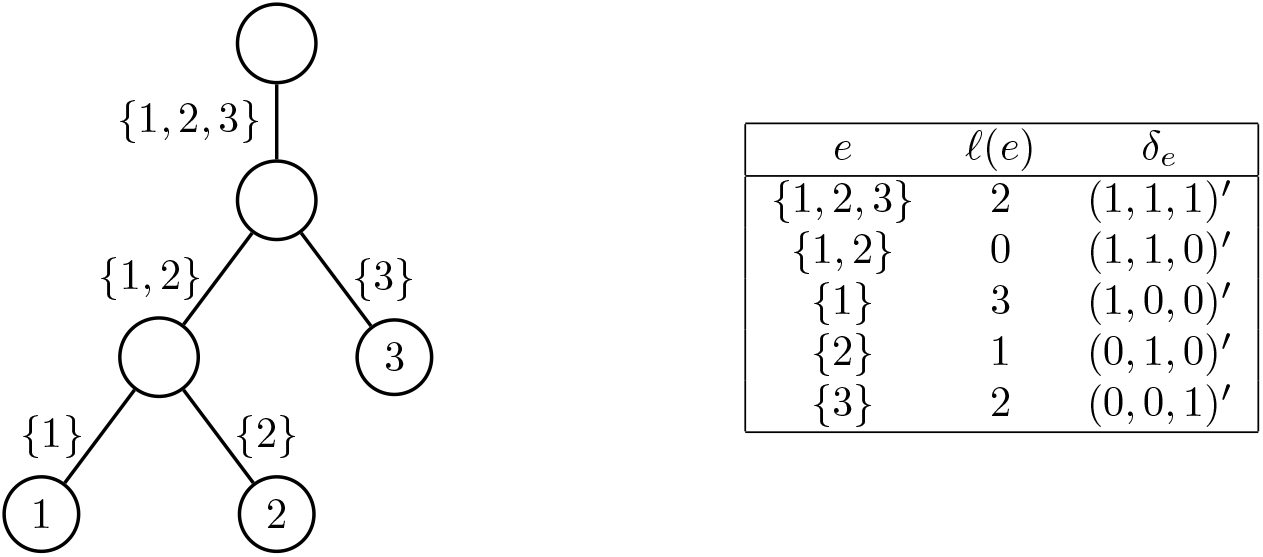
ORB-tree and strictly ultrametric matrix correspondence. The matrix encoding of the tree is 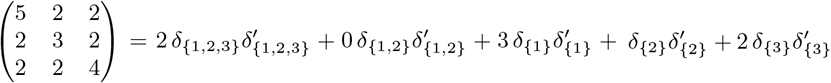.

Conversely, the matrix may be recovered from its ORB-tree as follows. For each edge *e* in the ORB-tree, let *δ*_*e*_ be the vector of dimension *n* with entries *δ*_*e*_(*i*) = 1 for *i* ∈ *e* and *δ*_*e*_(*i*) = 0 for *i* ∉ *e*. It follows from [44, Theorem 2.2] that

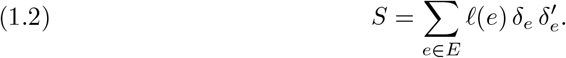

The representation of strictly ultrametric matrices as ORB-trees is therefore one-to-one. In fact, starting from an ORB-tree we may compute directly the entries of its associated matrix using that

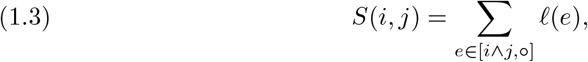

where the denotes the root of the tree. We may say therefore that the entries of a strictly ultrametric matrix are indexed by the leaves of its associated ORB-tree.

We call a matrix with entries such as (1.3) the covariance matrix of the ORB-tree. This terminology is borrowed from the ecology literature where matrices like this are commonly referred to as a tree-structured or phylogenetic covariance matrices [9]. In this setting, the leaves represent organisms, and the matrix entries denote a trait’s covariance between pairs of organisms. (The term of cophenetic matrix or cophenetic distance has also been used occasionally in the hierarchical clustering literature [50].)

Due to the identity in equation (1.3), strictly ultrametric matrices are usually fully dense. Nevertheless, precisely because of this identity, their entries contain much redundancy, suggesting they may be amenable to some form of compression. In this manuscript, we apply a change of bases—a discrete wavelet, in fact—with respect to which the covariance matrix of an ORB-tree often becomes sparse.

Wavelets are localized, wave-like functions developed to analyze non-stationary and noisy continuous signals. Traditional wavelets are defined only in Euclidean spaces and have been remarkably successful in identifying multiscale structures in signals and producing sparse representations of the same [39].

The Haar wavelet is among the oldest and involves averaging a signal locally at different time or space scales [23]. Recently, the authors of [21] extended it past continuous signals introducing the Haar-like wavelet. This new, discrete, wavelet is designed for the multiscale analysis of discrete datasets equipped with a partition tree—a hierarchical structure that clusters the data into smaller subsets recursively. Due to the organization of such datasets into different tree levels (i.e., scales) and clusters (i.e., localizations), Haar-like wavelets may identify meaningful patterns in data that would be impossible to detect otherwise—especially in noisy high dimensional datasets.

This paper exploits the equivalence between strictly ultrametric matrices and ORB-trees to sparsify the former via a change of basis. This basis is composed of the so-called Haar-like wavelets of the associated ORB-trees. The sparsification achieved by these wavelets can be substantial in large, strictly ultrametric matrices, which would otherwise be inaccessible due to their size. These sparse representations can be valuable in phylogenetic applications [47], network inference [30], and hierarchical clustering problems [50] as their models often rely on tree-structured covariance matrices.

### 1.1. Paper organization

In Section 2 we specialize the Haar-like basis from [21] and give a geometric interpretation of its action on ORB-trees. This results in a closed form expression for the transformed ultrametric matrix that can be computed efficiently—without having to pre-compute the matrix from the tree. In Section 3, we present conditions under which the Haar-like basis can be used to sparsify large, strictly ultrametric matrices, and show that the basis can substantially sparsify most large random ORB-tree’s covariance matrices. Following in Section 4, we detail the case in which the Haar-like basis diagonalizes a strictly ultrametric matrix and provide examples of such for well-known tree topologies. And, in Section 5, we show that the Haar-like basis can be used sometimes to estimate eigenvalues of ORB-tree’s covariance matrices.

Finally, Section 6 is devoted to an extensive proof of concept of our methods in metagenomics (i.e., the study of microbial environments based on genetic material extracted directly from them). Specifically, we sparify the covariance matrix associated with microbiologists’ Tree of Life. The significant sparsification achieved by our methods motivates introducing a new but wavelet-based phylogenetic (*β*-diversity) distance, corresponding to a multiscale analysis of organism abundances in microbial environments. This new distance gives remarkably similar results to other well-known metrics on a previously studied dataset. However, unlike the established metrics, it can also determine the splits in the Tree responsible for the observed microbial compositions and quantify their respective importance.

### 1.2. General notation and terminology

For real-vectors *x* = (*x*_*i*_)_1≤*i*≤*k*_ and *y* = (*y*_*i*_)_1≤*i*≤*k*_ of dimension *k*, let 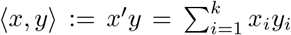 and 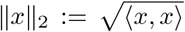. Also, let ⟦·⟧ denote the indicator function of the proposition within the parentheses.

In our context, trees are finite undirected connected graphs without cycles.

In what remains of this manuscript, *T* denotes an ORB-tree with *n* leaves and branch length function *ℓ* : *E* → ℝ. We denote the vertex and edge set of *T* as *V* and *E*, respectively. The root of *T* is denoted as ○. The set of internal nodes of *T* is denoted as *I*, whereas its set of leaves is denoted as *L*. By definition, ○ ∈*I* and *I* and *L* partition *V*. From the definition of ORB-tree it also follows that |*L*| = |*I*| = *n*, hence |*V*| = 2*n*. Note that |*E*| = |*V*| − 1 because *T* is a tree. We define |*T*| := |*V*|. We use this later notation when we want to emphasize a direct relationship with the ORB-tree.

For *i, j* ∈ *V*, a path of length *l* between *i* and *j* is a sequence *v*_0_, …, *v*_*l*_ ∈ *V* such that *v*_0_ = *i, v*_*l*_ = *j*, and {*v*_*k*_, *v*_*k*+1_} ∈ *E* for 0 ≤ *k < l*. Unless otherwise stated, we write [*i, j*] to denote the set of edges in the shortest path in *T* between *i* and *j*. This path is unique because *T* has no cycles. The depth of *i*, denoted depth(*i*), is defined as |[*i*, ○]| i.e. the number of edges that connect *i* with the root. We say that *i* is an ancestor of *j*, or alternatively *j* is a descendent of *i*, when *i* ∈[○, *j*]. In particular, every node is an ancestor and a descendant from itself. Further, (*i* ^ *j*) denotes the so-called least-common ancestor to *i* and *j*. This is the *v* ∈ *V* that maximizes |[*v*, ○]|, among all the nodes that are ancestors to both *i* and *j*.

We define

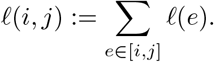

In addition, for *J* ⊂ *L* and *i* ∈ *V*, define *ℓ*(*J, i*) as the column vector of dimension |*J*| with entries *ℓ*(*j, i*), for *j* ∈ *J*. *ℓ*(*i, J*) is the transpose of *ℓ*(*J, i*).

For each *i* ∈ *V, T* (*i*) denotes the subtree of *T* rooted at *i*. In particular, the vertex set of *T* (*i*) is the subset of nodes in *T* that descend from *i*, and its edge set is the subset of edges that connect two descendants of *i. L*(*i*) denotes the leaf set of *T* (*i*). Likewise, for each *e* = *i, j E*, if *i* is a node closest to the root than *j, T* (*e*) and *L*(*e*) denote *T* (*i*) and *L*(*i*), respectively.

## 2. Haar-like basis of ORB-trees

In this section we specialize the concept of Haar-like basis given in [21] to our setting of ORB-trees. The key new result in this section is an expression for the change of basis of the covariance matrix (i.e. the strictly ultrametric matrix associated with the tree) into the Haar-like basis. The simplicity of this expression is somewhat unexpected because the covariance matrix is determined by the topology of the tree and its branch length function, whereas the basis is solely determined by the tree’s topology.

To construct the Haar-like wavelets, it is convenient to represent the nodes in *I*\ {*o*} momentarily as binary strings. With this convention, the (only) child of the root is denoted as *ε*—the so-called empty string. Further, the children of each node *v* ∈ *I*\ {*o*} are *v*0 (i.e. the string *v* with the character zero appended at the end) and *v*1 (i.e. *v* with the character one appended at the end).

### Definition 2.1 (Specialization from [21]).

*The Haar-like basis associated with T is the set of transformations {φ*_*v*_*}*_*v*∈*I*_ *defined as follows:*

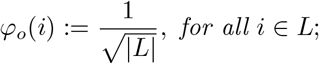

*and, for each v* ∈ *I with v* ≠ *o:*

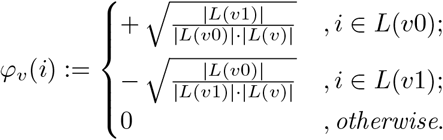

*The Haar-like matrix associated with T is the matrix* Φ *with columns φ*_*v*_, *v* ∈ *I*.

To fix ideas see Figure 2.

**Fig. 2:**
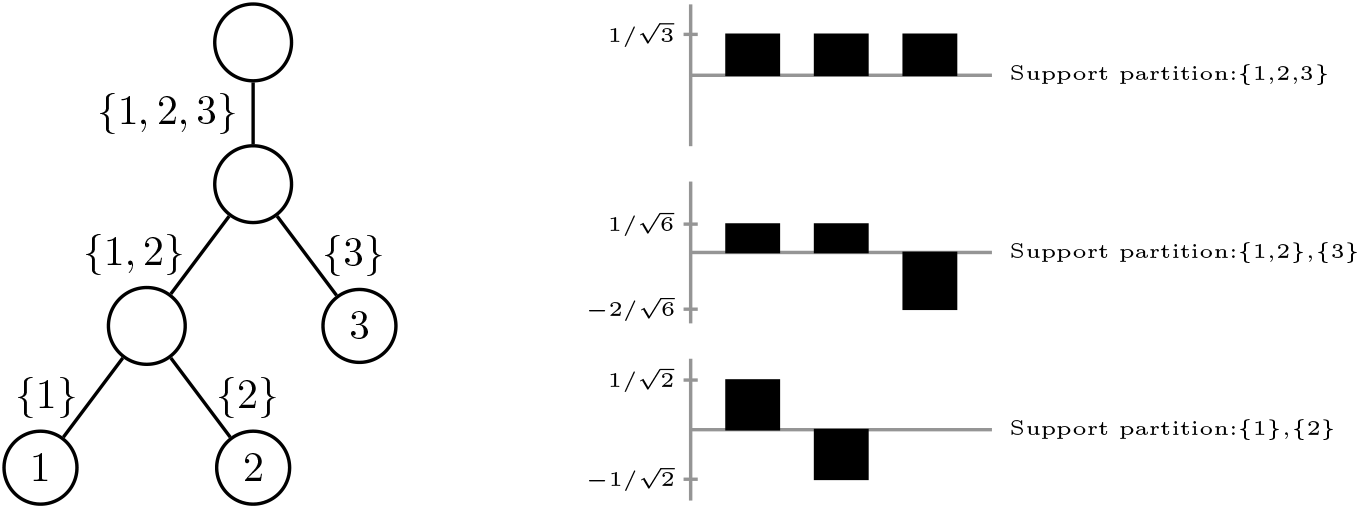
Visualization of the Haar-like Wavelet basis associated with an ORB-tree. Left: ORB-tree with leaves 1, 2, 3. Edges are labeled by the subsets of leaves that descend from them. Right: Haar-like basis associated with the ORB-tree on the left.

We remark that, in the context of compositional data analysis, the Haar-like basis is equivalent to the orthonormal basis used in the isometric log ratio (ILR) transform [15]. We revisit the implications of this connection in Section 6.

The terminology of basis in Definition 2.1 is justified by the fact that

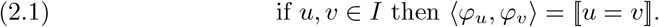

(From now on, ⟦·⟧ denotes the indicator function of the proposition within.) In particular, {*φ*_*v*_}_*v*∈*I*_ is an orthonormal basis of ℝ ^|*L*|^. (See the Appendix for a self-contained justification of the orthonormality of the Haar-like basis.) Note that the Haar-like matrix Φ has its rows indexed by *L* and its columns indexed by *I*. Hence, since |*L*| = |*I*|, Φ is a square matrix, an orthonormal one.

Clearly, for each *v* ∈ *I, φ*_*v*_ has *L*(*v*) as its support. constant, the orthogonality property implies for *v* ≠ *o* that *∑*_*i*∈*L*_ *φ*_*v*_(*i*) = 0. These two properties are essential for our arguments onwards.

The following definition is useful to understand the relationship between the Haar-like basis of an ORB-tree and its associated covariance matrix.

### Definition 2.2.

*The trace branch length of T is the function ℓ*\* : *E* → [0, ∞) *defined as ℓ*\*(*e*) := |*L*(*e*)| *ℓ*(*e*), *for each e* ∈ *E*.

### Theorem 2.3.

*If v* ∈ *I then S φ*_*v*_ = diag(*ℓ*\*(*L, v*)) *φ*_*v*_.

*Proof*. Consider *v* ∈ *I* and *j* ∈ *L*. If *j* ∉ *L*(*v*) then (*i* ∧ *j*) = (*v* ∧ *j*), hence

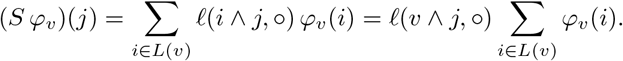

But, if *v* = ○ then *ℓ*(*v* ∧ *j*, ○) = 0. Instead, if *v* ≠ ○ then *∑*_*i* ∈*L*(*v*)_ *φ*_*v*_(*i*) = 0. In either case: (*S φ*_*v*_)(*j*) = 0. This shows the lemma for *j* ∉ *L*(*v*) because the entry associated with *j* in diag(*ℓ*\*(*L, v*)) *φ*_*v*_ is *ℓ*\*(*v, j*) · *φ*_*v*_(*j*), and the support of *φ*_*v*_ is *L*(*v*).

Next suppose that *j* ∈ *L*(*v*). Then

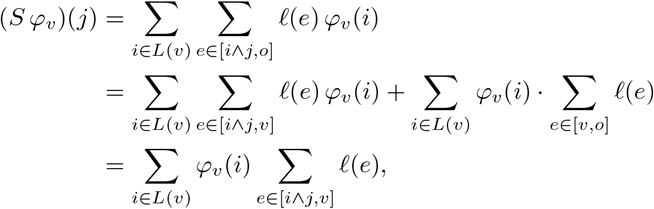

where for the last identity we have used that *∑*_*i*∈*L*(*v*)_ *φ*_*v*_(*i*) = 0 if *v* ≠ ○, and ∑_*e*∈[*v,o*]_ *ℓ*(*e*) = 0 if *v* = ○. But note that if *i* ∈ *L*(*v*) is such that (*i* ∧ *j*) = *v* then *∑*_*e*∈[*i*∧*j,v*]_ *ℓ*(*e*) = 0. Instead, if (*i* ∧ *j*) ≠ *v* then *φ*_*v*_(*i*) = *φ*_*v*_(*j*). As a result

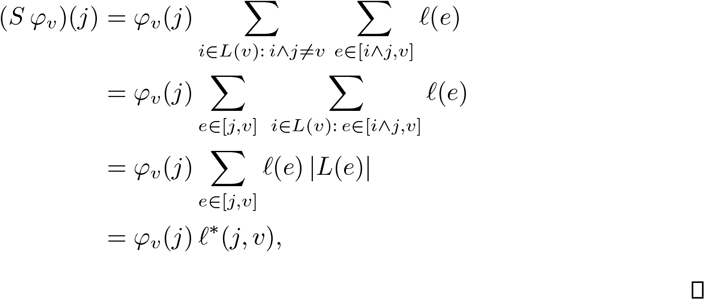

which shows the result.

It follows from the theorem that for each *u, v* ∈ *I*:

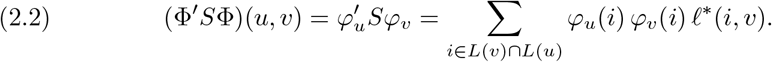

The importance of the diagonal of Φ′*S*Φ in the discussion ahead, motivates to define for *v* ∈ *I* the quantities

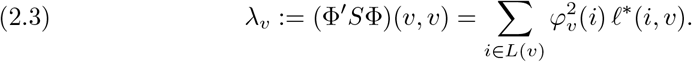

For *v* ∈ *I*, because *φ*_*v*_ has *L*(*v*) as its support and ∥*φ*_*v*_∥ _2_ = 1, *λ*_*v*_ is a weighted average of the trace branch length between each leaf in *L*(*v*) and *v*. In particular, since *L*(*u*) ⊃ *L*(*v*) when *u* is an ancestor of *v*, the closer the internal node *v* is to the root, the more terms are averaged. (This emulates the averaging at different scales that the standard Haar wavelet transform does to a continuous signal.) Furthermore, since *ℓ*\*(*e*) = *ℓ*(*e*) *>* 0 when *e* joins a leave with its parent, *λ*_*v*_ *>* 0.

On the other hand, the identity in (2.2) implies that (Φ′*S*Φ)(*u, v*) = 0 when *u, v* ∈ *I* are such that *L*(*u*) ⋂ *L*(*v*) = ∅. This suggests that the Haar-like matrix can be used to sparsify the covariance matrix of the ORB-tree. The following result is critical to assess how effective this sparsification is in practice.

### Lemma 2.4.

*For all u, v* ∈ *V, L*(*u*) ⋂ *L*(*v*) ≠ ∅ *if and only if u is an ancestor of v or vice versa*.

*Proof*. If *u* is an ancestor of *v* then *L*(*v*) ⊂ *L*(*u*); in particular, *L*(*u*) ⋂ *L*(*v*) = *L*(*v*) ≠ ∅. The same conclusion applies if *v* is an ancestor of *u*. Conversely, suppose that *L*(*u*) ⋂ *L*(*v*) ≠ ∅. Without loss of generality assume that *u* ≠ *v*. From the hypothesis, there is *w* ∈ *L* that descends from both *u* and *v*. But, since there is a unique path from *w* to ○, *u* and *v* must be both in this path; in particular, either *u* is an ancestor of *v* or vice versa.

### 2.1. Fast Sparsification Algorithm

A non-trivial challenge to storing and manipulating large strictly ultrametric matrices is that they are almost always fully dense in practice. In many applications, particularly metagenomics, the ORB-trees associated with such matrices are assumed to be known in advance. This allows us to sparsify these matrices without computing them, or even storing them in computer memory. It also allows us to anticipate which entries may remain nonzero after sparsification. In fact, due to equation (2.2) and Lemma 2.4, all that is required to sparsify these matrices from their ORB-tree is to precompute the leaves that descend from each internal node (i.e., the sets *L*(*v*), with *v* ∈ *I*) and the trace branch length between them (Definition 2.2). This can be achieved with two postorder traversals of the ORB-tree. We convey these ideas in the following pseudo-code (Algorithm 2.1), which is fully coded and available on GitHub.

#### Algorithm 2.1

Phylogenetic covariance matrix sparsification

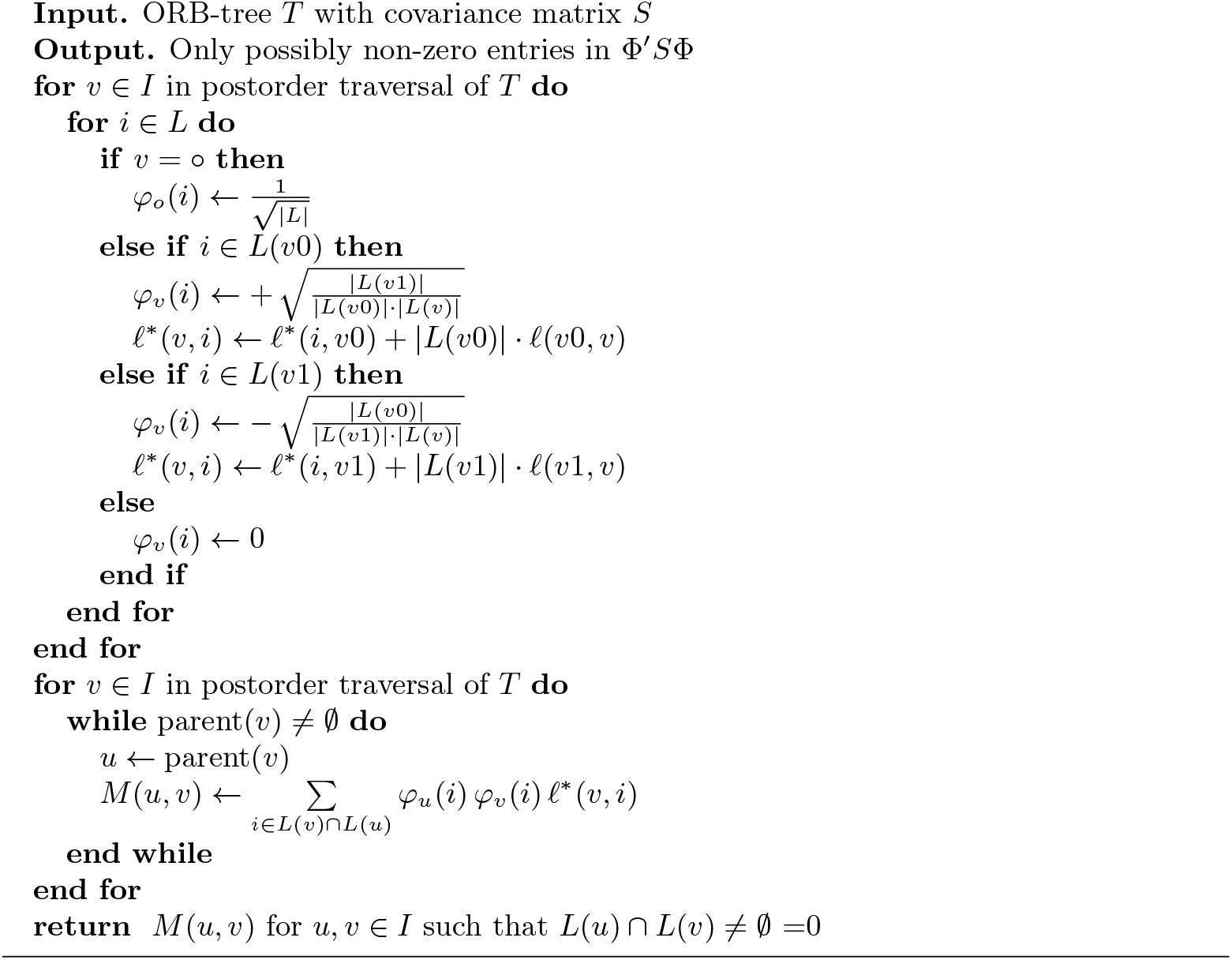

## 3. Sparisfication of Covariance Matrices of ORB-trees

In this section we quantify how much of the covariance matrix of an ORB-tree can be sparsified by its Haar-like matrix. To state our main result we require the following definitions.

### Definition 3.1.

*Recall that* |*T* | *denotes the total number of nodes in T. The average subtree size of T is the quantity* avg 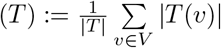.

### Definition 3.2.

*The internal and external path lengths of T are the quantities defined as* IPL(*T*) := *∑*_*v*∈*I*_ *depth*(*v*) *and* EPL(*T*) := *∑*_*v*∈*L*_ *depth*(*v*), *respectively [51]. The total path length of T is the quantity* TPL(*T*) := IPL(*T*) + EPL(*T*).

We note the relationship:

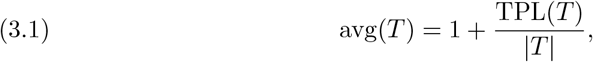

because

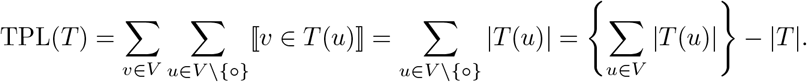

### Definition 3.3.

*The interior of T is the tree* 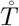 *obtained by trimming the leaves of T*.

Clearly, 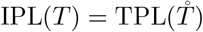.

As mentioned earlier, the identity in (2.2) guarantees that some entries of Φ′*S*Φ vanish. The following result gives a lower bound for the number of such entries. This bound is independent of the branch lengths and depends—only—on the tree topology.

### Theorem 3.4.

*If ζ denotes the fraction of vanishing entries in* Φ^*′*^*S*Φ *then*

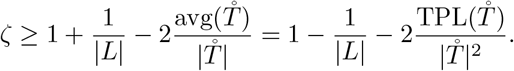

*Proof*. Recall that |*I*| = |*L*| and, for *ν* ∈ *I*, the support of *ϕ*_*v*_ is *L*(*ν*). Hence, from the identity in (2.2), (Φ^*′*^*S*Φ)(*u, ν*) = 0 when *u, ν* ∈ *I* and *L*(*u*) ∩ *L*(*ν*) = ∅. As a result, using that *L*(*u*) *≠* ∅ when *u* ∈ *I*, and Lemma 2.4, we obtain that

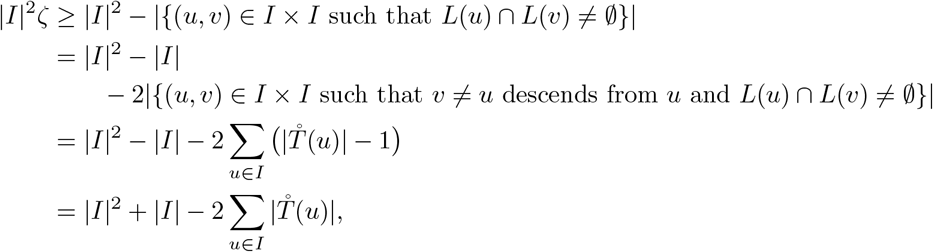

Since 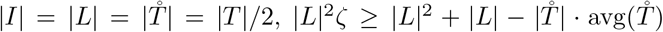, which shows the inequality in the theorem. The alternative lower-bound for *ζ* follows by applying the identity in equation (3.1) to 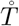, completing the proof of the theorem.

It follows from the first lemma in [51, Section 6.4] that for an ORB-Tree T, EPL(*T*) − IPL(*T*) = 2 |*I*| − 1, which together with the previous theorem let us conclude the following asymptotic result.

### Corollary 3.5.

*If either* 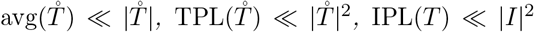, *or* EPL(*T*) *≪*|*L*|2 *as* |*T* | → ∞, *then ζ* = 1 − *o*(1).

In other words, if *T* grows so that either of the asymptotic inequalities in the above corollary applies, then an asymptotically negligible fraction of the off-diagonal entries in Φ^′^*S*Φ will be non-zero.

The last asymptotic condition in the corollary (i.e., that EPL(*T*) *≪*|*L*|^2^) is of relevance in phylogenetic studies. In that context, the external path length of a tree is called its Sackin’s index [3, 10, 32]. This index is used as a measure of the imbalance of phylogenetic trees. In particular, since phylogenetic trees are neither too balanced nor too imbalanced [2, 4]; Haar-like bases should be rather effective in sparsifying covariance matrices of phylogenetic trees in practice. We come back to this point in Section 6.

### 3.1. Covariance matrices of maximally balanced ORB-trees

In our context, the following definition gives the most balanced topology among the ORB-trees.

*A perfect binary tree is an ORB-tree in which all leaves have the same depth*.

To fix ideas see Figure 3.

**Fig. 3:**
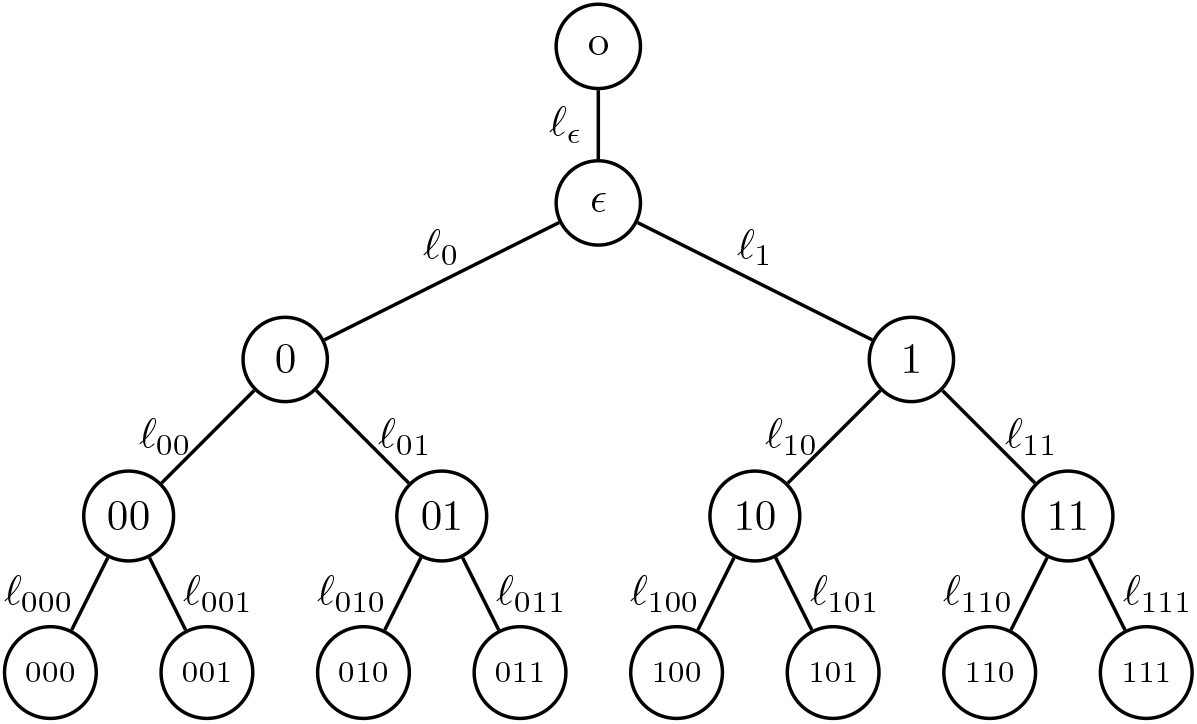
Visualization of a perfect binary tree of height 4. Such tree is trace-balanced if and only if *ℓ*_*α*_ = *ℓ*_*β*_ for each pair of binary strings *α* and *β* of the same length. If all these lengths are strictly positive, the eigenvalues of its covariance matrix are *ℓ*_000_ (multiplicity 4), *ℓ*_000_ + 2 *ℓ*_00_ (multiplicity 2), *ℓ*_000_ + 2 *ℓ*_00_ + 4 *ℓ*_0_ (multiplicity 1), and *ℓ*_000_ + 2 *ℓ*_00_ + 4 *ℓ*_0_ + 8 *ℓ* (multiplicity 1). Otherwise, some multiplicities need to be added up.

Let *T* be a perfect binary tree of height (*h* + 1). In particular, |*L* |= 2^*h*^ and |*V*| = 2^*h*+1^. At level *k* ≥ 1, *T* contains 2^*k*−1^ nodes, each of which is the root of a perfect binary tree of height (*h* − *k*). Since a perfect binary tree of height *h* contains (2^*h*+1^ − 1) nodes, and the interior of a perfect binary tree of height (*h* + 1) is a perfect binary tree of height *h*:

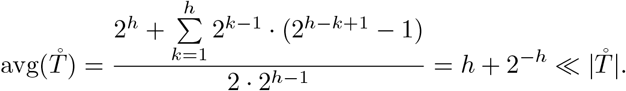

In particular, due to Theorem 3.5, we can conclude that the Haar-like matrix can be used to asymptotically annihilate (via a similarity transformation) the off-diagonal entries of the covariance matrix of a perfect binary tree as its height tends to infinity.

### 3.2. Covariance matrices of maximally imbalanced ORB-trees

The following definition provides what we may regard as the most unbalanced topology among the ORB-trees.

#### Definition 3.7 (Binary Caterpillar Trees).

*The binary caterpillar tree of height h* ≥ 1 *is the ORB-tree with nodes ○*, 1, …, *h*, 1^*′*^, …, (*h* − 1)^*′*^ *and edges of the form {○*, 1*}, {i, i* + 1*}, for i* = 1, …, (*h* − 1), *and {j, j*^′^*} for j* = 1, …, (*h* − 1).

To fix ideas see Figure 4.

**Fig. 4:**
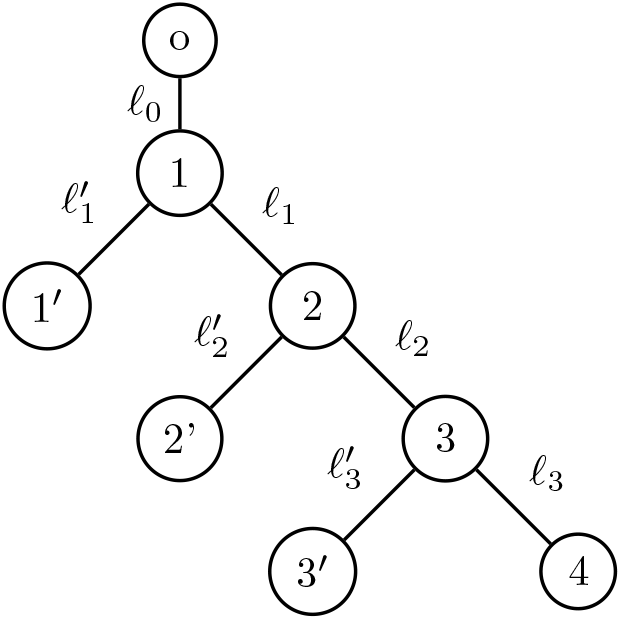
Visualization of a binary caterpillar tree of height 4. This tree is trace-balanced if and only if 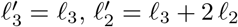, and 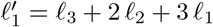. In such case, if *ℓ*_3_, *ℓ*_2_, *ℓ*_1_, *ℓ*_0_ *>* 0 then its covariance matrix spectrum is {ℓ_3_, *ℓ*_3_ + 2 *ℓ*_2_, *ℓ*_3_ + 2 *ℓ*_2_ + *ℓ*_1_, *ℓ*_3_ + 2 *ℓ*_2_ + 3 *ℓ*_1_ + 4 *ℓ*_0_}, and each eigenvalue is simple.

Let *T* be a binary Caterpillar tree of height *h*. In particular, |*V* |= 2*h*, |*L*| = *h*, and each node in level *k* ≥ 1 is the root of a snake binary subtree of height (*h* − *k*). Therefore, each internal node has as children one leaf node and one internal node that is the root of a binary caterpillar subtree. Furthermore, the subgraph of internal nodes is a path. As a result:

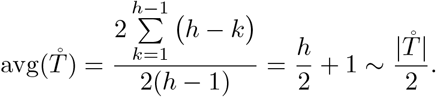

Hence, the lower-bound provided by Theorem 3.4 is trivial, and we cannot guarantee that the Haar-like matrix associated with a sizeable binary caterpillar tree annihilates its off-diagonal entries in any significant way.

### 3.3. Covariance matrices of large random ORB-trees

Perfect binary trees and caterpillar trees are opposite extremes of how balanced (or imbalanced) ORB-trees can be. It is therefore unclear how much sparsification the Haar-like matrix of a large but generic ORB-tree can induce on its covariance matrix. To address this issue we consider a natural ensemble of random ORB-trees

In what follows, 𝕋 denotes a uniformly at random ORB-tree with |*I*| internal nodes. Such trees may be generated using the Catalan distribution [51, Section 6.7].

This probability model produces full binary trees (i.e. trees in which each node has 0 or 2 children) with a given number of internal nodes; which we may turn into an ORB-tree by appending their root to a new one.

Let 𝕊 denote the covariance matrix of 𝕋, and *ζ* the number of zeroes in the random matrix Φ^*′*^𝕊Φ, where Φ is the Haar-like matrix associated with 𝕋. It turns out that the mean and variance of the internal path length of 𝕋 are given by

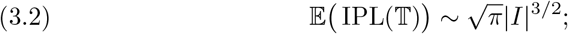

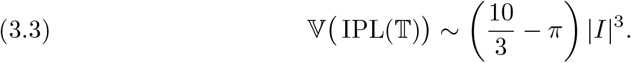

The identity in equation (3.2) follows from [18, Proposition VII.3.]. The identity in (3.3) may be regarded a refinement of [18, Note VII.12].

As the following result implies, the Haar-like basis of most large ORB-trees should be highly effective in sparsifying their covariance matrix.

#### Corollary 3.8.

*If 𝕋 is a uniformly at random ORB-tree with* |*I*| *internal nodes then ζ* = 1 − *o*(1) *with overwhelmingly high probability, as* |*I*| → ∞.

*Proof*. Let *t >* 0. Let *μ* and *σ*^2^ denote the mean and variance of IPL(𝕋), respectively. Due to Cantelli’s inequality (a one sided version of the well-known Chebyshev’s inequality): ℙ IPL(𝕋) ≥ *μ* + *tσ* ≤ (1 + *t*^2^)^−1^. But (*μ* + *tσ*) = Ω (*t* |*I* ^3′2^) because of equations (3.2)-(3.3). In particular, there is a constant *c >* 0 such that

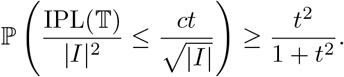

So, if *t* so that 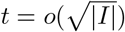 then 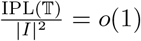 with a probability converging to one as |*I*| → ∞. The result now follows from Corollary 3.5.

## 4. Spectrum of Covariance Matrices of Trace-balanced ORB-trees

While Theorem 3.4 guarantees that some entries in Φ^*′*^*S*Φ vanish—regardless of branch lengths, additional constraints on the latter can lead to further sparsification. In this section, we identify the class of ORB-trees whose Haar-like basis fully sparsifies (i.e, diagonalizes) their associated strictly ultrametric matrix.

### Definition 4.1.

*T is called trace-balanced at a node ν when, for all i, j* ∈ *L*(*ν*), *f*\*(*i, ν*) = *f*\*(*j, ν*). *T is called trace-balanced when it is trace-balanced at each ν* ∈ *I \ {○}*.

We note that a tree is always trace-balanced at a leaf. Also, if an ORB-tree is trace-balanced at the child of its root then it is also trace-balanced at the root. This is why the definition of trace-balanced trees only considers nodes in *I* \ {*o*}. See Figures 3-4 for depictions of trace-balanced trees.

The following two results show the relevance of the above definition in terms of the eigenvalues of the ultrametric matrix associated with an ORB-tree.

### Lemma 4.2.

*If ν*∈ *I then ϕ*_*v*_ *is an eigenvector of S if and only if T is trace-balanced at ν, in which case the eigenvalue associated with ϕ*_*ν*_ *is f*\*(*i, ν*), *for any i* ∈ *L*(*ν*).

*Proof*. Fix *ν* ∈ *I*. Due to Theorem 2.3, (*Sϕ*_*ν*_)(*i*) = *f*\*(*i, ν*) *ϕ*_*ν*_(*i*), for each *i* ∈ *L*. This shows the lemma because *ϕ*_*ν*_(*i*) *>* 0 if and only if *i* ∈ *L*(*ν*).

Since the covariance matrix *S* of *T* has dimensions |*L*|× |*L*|, but |*I*| = |*L*| because *T* is an ORB-tree, the following result is immediate from the previous lemma.

### Corollary 4.3.

*The Haar-like basis of T diagonalizes its covariance matrix if and only if T is trace-balanced. In this case, the spectrum of S is*

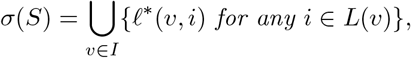

*and the multiplicity of f*\*(*ν, i*) *is {u* ∈ *I* : *f*\*(*ν, i*) = *f*\*(*u, j*), *for some j* ∈ *L*(*u*)*}*.

Our next corollary recovers some of the results in [48], which investigated perfect binary trees in the phylogenetic setting with the goal of predicting features of protein structure.

### Corollary 4.4.

*A perfect binary tree of height h is trace-balanced if and only if it has constant branch lengths at each level. In this case, if* 𝓁_*j*_ *denotes the common length of the edges that connect a node at depth j with another at depth* (*j* + 1), *the spectrum of the associated covariance matrix is*

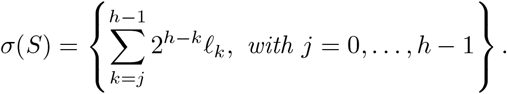

*Furthermore, the multiplicity of the eigenvalue* 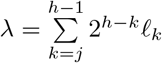 *is* max 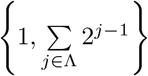

*where*

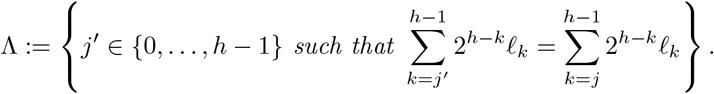

Next, consider the binary caterpillar tree from Definition 3.7. In particular, its internal and leaf set are *I* = {○, 1, …, *h*−1*}* and *L* = {1^*′*^, …, (*h*−1)^*′*^, *h*}, respectively. Let 𝓁_0_ denote the branch length of {○, 1}, 𝓁_*i*_ the length of {*i, i*+1} for *i* = 1, …, (*h*−1), and 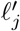 the branch length of {*j, j*^*′*^} for *j* = 1, …, (*h* − 1). Due to Corollary 4.3, we have the following result.

### Corollary 4.5.

*A Caterpillar tree of height h is trace-balanced if and only if*

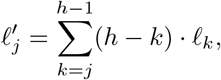

*for j* = 1, …, *h* − 1. *In this case, the eigenvalues of its covariance matrix are as follows, repeated according to their multiplicity:* 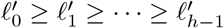, *where*

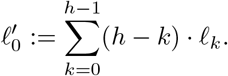

We finish this section with a result that characterizes all the possible spectrums of covariance matrices of trace-balanced trees. Because its proof is constructive, it can be used to form strictly ultrametric matrices with the desired spectrum and multiplicities. In particular, it is of consequence for the symmetric nonnegative inverse eigenvalue problem (SNIEP), which aims to classify the possible spectra of symmetric nonnegative matrices [54, 16, 13].

### Definition 4.6.

*In a tree T, a function f* : *I* → [0, ∞) *is called decreasing when, for all distinct u, ν*∈ *I, if u is an ancestor of ν then f* (*u*) ≥ *f* (*ν*). *In addition, f is called strictly positive at the fringe when f* (*u*) *>* 0 *whenever u is a parent of a leaf*.

### Corollary 4.7.

*In a trace-balanced ORB-tree T the function ν* −→ *f*\*(*ν, i*), *with ν* ∈ *I and any i* ∈ *L*(*ν*), *is decreasing, and strictly positive at the fringe. Conversely, given any ORB-tree topology T and decreasing function f* : *V* → [0, ∞) *that is strictly positive at the fringe, there is a branch length function f* : *E* → [0, ∞) *such that σ*(*S*) = *f* (*I*). *Furthermore, the multiplicity of λ* ∈ *σ*(*S*) *is* |*f* ^−1^(*{λ}*)|.

*Proof*. From the definitions of trace-balanced and ORB-tree, it is immediate that the transformation *ν* ∈ *I* −→ *ℓ*\*(*ν, i*), with *i* ∈ *L*(*ν*), is well-defined and strictly positive at the fringe of *T*. Also, *f* is decreasing because if *u* is an ancestor of *ν* then, for each *i* ∈ *L*(*ν*): *ℓ*\*(*u, i*) = *ℓ*\*(*u, ν*) + *ℓ*\*(*ν, i*) ≥ *ℓ*\*(*ν, i*). This shows the first statement in the corollary.

For the second statement consider an ORB-tree topology *T* = (*V, E*) and function *ℓ* : → *I* [0, ∞) that is both decreasing and strictly positive at the fringe. Due to Corollary 4.3, it suffices to show that there is a branch length function *ℓ* : → *E* [0, + ∞) such that *ℓ* (*ν*) = *ℓ*\*(*ν, i*), for all *i* ∈ *L*(*ν*). To do so, let *e* = {*u, ν}* ∈ *E* be so depth(*u*) *<* depth(*ν*). Define

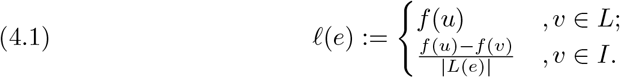

Observe that if *ν* ∈ *L* then |*L*(*e*)| = 1 so *f* (*u*) = *ℓ*\*(*e*). Further, *ℓ*(*e*) *>* 0 because *f* is strictly positive at the fringe of *T*. Instead, if *ν* ∈ *I* then *ℓ* (*u*) = *ℓ* (*ν*) + *ℓ*(*e*) · |*L*(*e*) | = *ℓ* (*ν*) + *ℓ*\*(*e*), and *ℓ*(*e*) ≥ 0 because *f* is decreasing. In particular, if we extend the domain of *f* to all of *V* defining *ℓ* (*ν*) := 0 for *ν L* then, for all *e* = {*u, ν*} ∈ *E* such that depth(*u*) *<* depth(*ν*): *ℓ* (*u*) = *ℓ* (*ν*) + *ℓ*\*(*e*). From this, a simple inductive argument on the difference *d* := depth(*ν*) − depth(*u*) *>* 0 shows that *f* (*u*) − *ℓ* (*ν*) = *ℓ*\*(*u, ν*); implying that *f* (*u*) = *ℓ*\*(*u, i*), for all *i* ∈ *L*(*u*), as claimed.

## 5. Spectrum Approximation in Roughly Trace-balanced ORB-trees

The ORB-tree representation of strictly ultrametric matrices offers new approaches to studying their spectrum. This is of interest for PCA, as well as domains such as structural biology [48] and metagenomics [45], where strictly ultrametric matrices emerge as covariance ones. This section examines how to approximate the spectrum of strictly ultrametric matrices associated with possibly non-trace-balanced ORB-trees. We have seen that the Haar-like wavelet associated with an internal node of an ORB-tree is an eigenvector of its covariance matrix if and only if the node is trace-balanced (Lemma 4.2). We have also seen that the Haar-like matrix of an ORB-tree can sometimes sparsify its covariance matrix significantly (Theorem 3.4). These facts suggest that the diagonal entry in Φ^*𠌣*^*S*Φ associated with an “approximately” trace-balanced internal node should be near the spectrum of *S*. Next, we formalize this intuition by quantifying what suffices for an internal node to be approximately trace-balanced. We stress that when our upper bound for the approximation error is small, the approximate eigenvalue can be computed efficiently using equation (2.3).

In what follows, for a given function *x* : *L* → *ℝ* and non-empty *J*⊂ *L*, we define the mean value and variance of average value and variance of *x* over *J* naturally as the quantities:

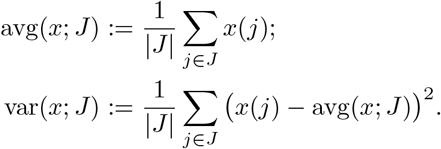

In addition, for each *ν* ∈ *V*, let parent(*ν*) denote the parent of node *ν* in *T*. Define

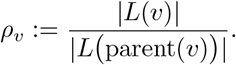

The following simple result aids in formalizing the intuition mentioned earlier.

Lemma 5.1. *If A is a symmetric matrix of dimensions n* × *n then, for all λ* ∈ ℝ:

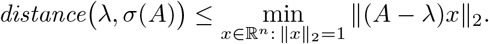

Next, we provide a sufficient condition for *λ*_*v*_, with *ν* ∈ *I*, to be a good approximation of an eigenvalue of *S*. We also quantify explicitly the cosine between *Sφ*_*v*_ and *λvφ*_*v*_ to assess how close *φ*_*v*_ is to be an eigenvector of *S*.

In the following result we use the notation: *¬*0 = 1 and *¬*1 = 0.

### Theorem 5.2.

*If ν* ∈ *I then*

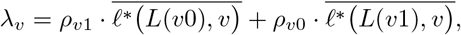

*and*

*distance*(*λv, σ*(*S*))

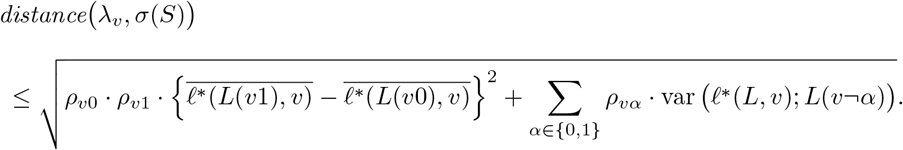

*Furthermore*,

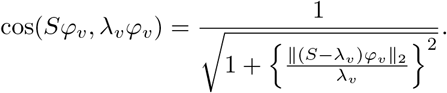

*Proof*. Fix *ν*∈ *I*. To make the *λ*_*v*_ more explicit, observe that if *x* : *L*→ *ℝ* is a function (or vector) then

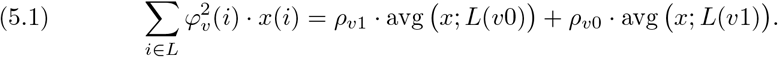

In particular, due to Theorem 2.3:

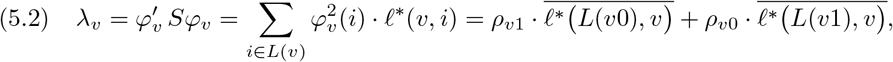

which shows the first identity in the theorem.

On the other hand, Lemma 5.1 implies that

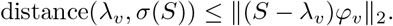

(But, from Theorem 2.3, we also have for *i* ∈ *L* that (*Sφ* _*v*_ − *λ*v *φv*)(*i*) = *φ* _*v*_ (*i*) ·(*ℓ*\*(*ν, i*) − *λ*_*v*_). As a result

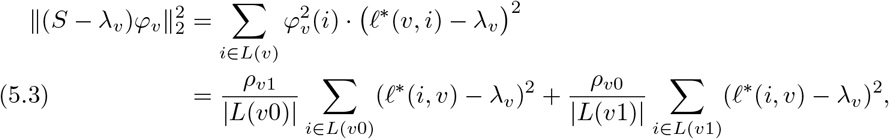

where for the last identity we have used the equation (5.1). To complete the proof of the theorem note that (*ρ* _*v*0_ *+ ρ* _*v*1_) = 1. In particular, from the identity in equation (5.2), we may rewrite

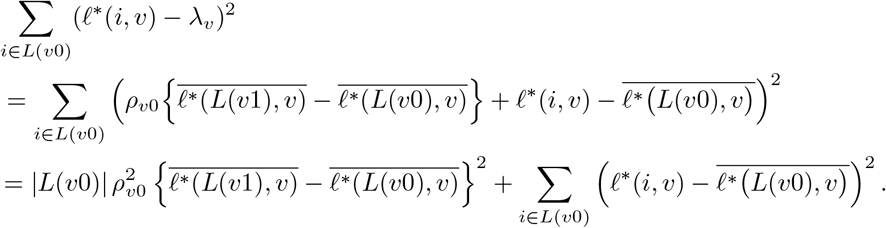

Namely

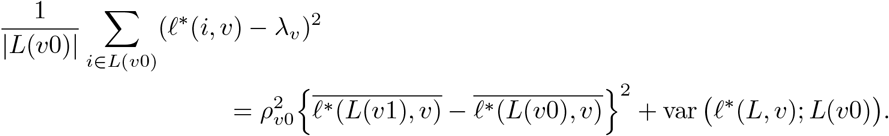

Similarly,

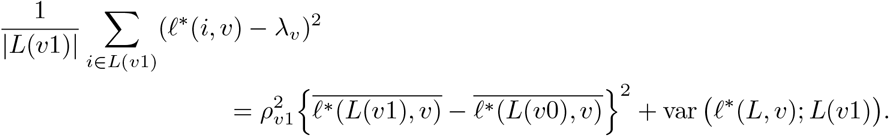

The second identity in the theorem is now a direct consequence of (5.3) and the last two identities.

Finally, again due to Theorem 2.3, we find that

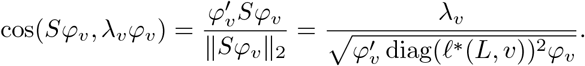

But, similarly as we argued before

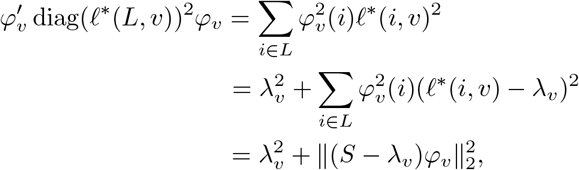

which implies that

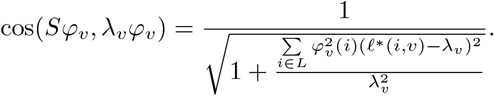

The third identity in the theorem follows now from equation (5.3).

It follows that *Sφ* = *λvφv* if and only if 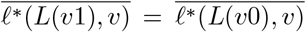 and also var (*ℓ*\*(*L, ν*); *L*(*ν*0)) = var (*ℓ*\*(*L, ν*)); *L*(*ν*1)) = 0. But these conditions are precisely equivalent to having the ORB-tree trace-balanced at *ν*. Lemma 4.2 may be therefore regarded a corollary of Theorem 5.2.

## 6. New Insights into the Microbial Tree of Life

In this section we apply our results to a phylogenetic covariance matrix associated with a standard reference phylogeny. The significant sparsification of this matrix opens the door for otherwise impossible tasks related to this model, such as computing the spectrum or inverse of its covariance matrix*—*standard tasks in phylogenetic comparative methods. In addition to the sparsification, the spectral approximation properties motivate the a new wavelet based metric used to compare microbial environments.

Many methods in microbiology rely on a phylogenetic tree relating microorganisms. At the microbial level, however, the notions of genus or species are ill-defined because microorganisms do not interbreed. So microbes’ taxonomy and phylogeny are often based on the so-called 16S ribosomal RNA (16S rRNA) gene. This gene is present in all known single cell organisms and can therefore be used as a phylogenetic marker. An operational taxonomic unit (OTU) is a cluster of these markers defined by some least level of DNA sequence similarity among its (highly) conserved regions. Greengenes is a standarized database based on the 16S rRNA marker. It has been a standard reference in microbial studies, particularly metagenomics, and is the default option in QIITA [22]*—*a widely used open-source management platform for microbial analyses. Greengenes phylogenetic trees are built using FastTree [46] and their associated taxonomies are assigned using tax2tree [41]. Trees are typically stored in the newick format [17], which encodes their topology and branch lengths, and visualized using software such as FigTree [1] or the ETE Toolkit [26].

Figure 5 displays the Greengenes tree when OTUs are thresholded at a 97*%* sequence similarity*—*the average similarity of macro-organisms’ DNA in the same species. The tree represents the inferred evolutionary history of modern day micro-organisms from common ancestors. Its root is at the center of the circular layout, and each OTU is associated with a single leaf in the tree and vice versa. Branch lengths are a proxy of evolutionary time such as the estimated expected number of mutations per nucleotide site [25], and interior nodes (called splits) are inferred speciation events that have led to the present-day microorganisms in the database.

**Fig. 5:**
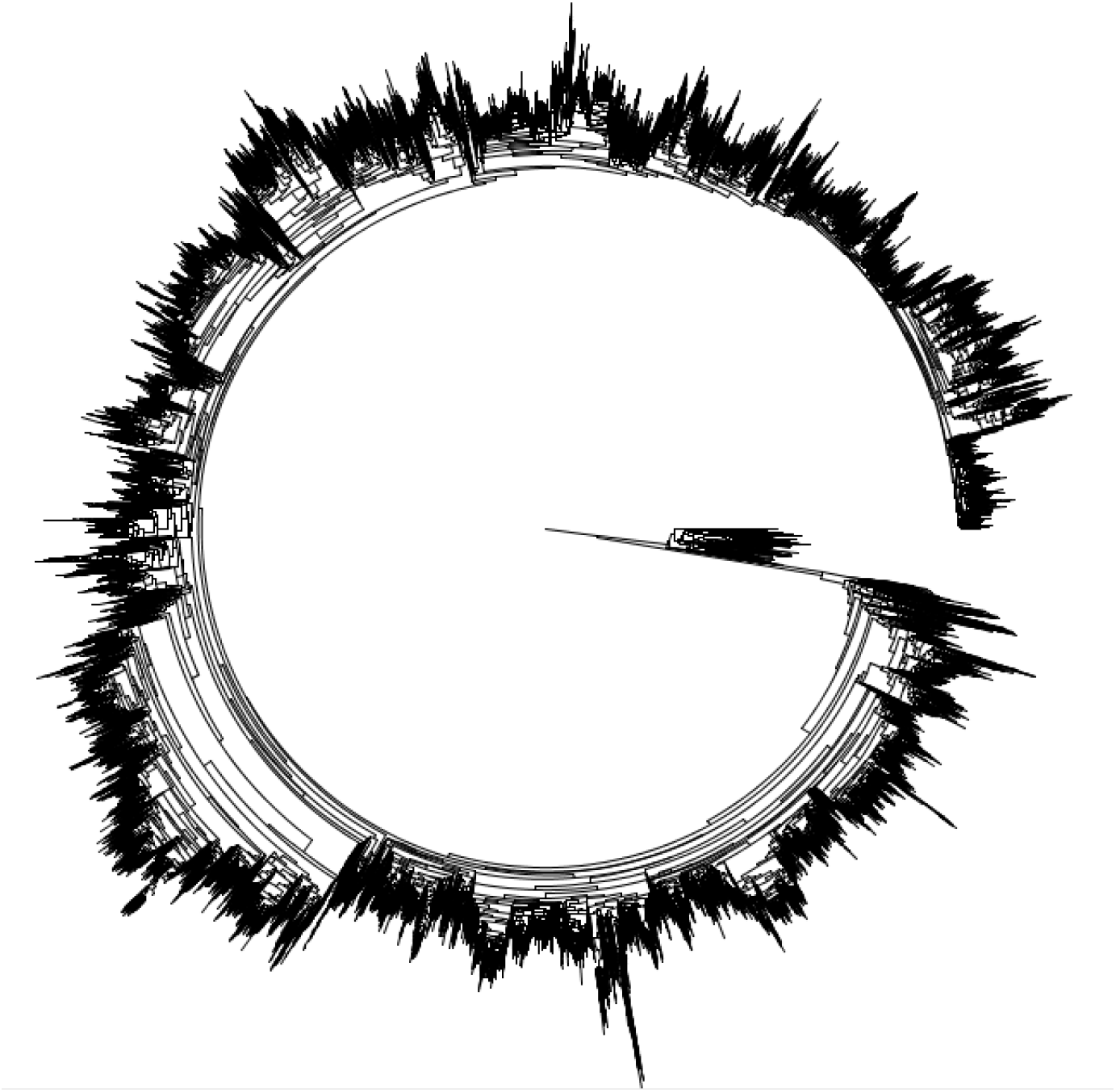
Circular layout of the 97*%* Greengenes tree. The tree has 99,322 leaves, 198,642 edges, and height (i.e. maximal leaf depth) 107. The average branch length is 1.42 × 10^−2^ units, with lengths varying between 1.5 × 10^−4^ and 1.0.

A fundamental problem in microbiology is to link environmental factors (such as acidity, light, nutrients, salinity, temperature, etc) with microbial composition. A valuable tool for this has been the concept of *β*-diversity (i.e., a measure of differences between microbial composition across different environments). Early approaches [6, 28] ignored the evolutionary relationships between microorganisms when comparing environments. Nonetheless, one would expect microbes with a shared evolutionary history to similarly thrive or struggle in similar environments. Phylogenetic informed metrics were introduced precisely to convey this idea. These metrics require a phylogenetic tree relating the microorganisms observed in samples from all the environments under study. We emphasize that the construction and selection of these trees are outside the scope of this paper; as is typical in many metagenomics studies, we work with a pre-computed tree. Among other more recent phylogentic trees such as SILVA [49] and WoL [60], Greengenes has been a common choice of representative phylogeny. So, we base our application on the latter*—*though our methods could be applied to any reference phylogeny.

Double Principal Coordinate Analysis (DPCoA) [45] is a phylogenetically informed *β*-diversity metric between pairs of microbial environments, which provides similar insights [20] to other more recent though more widely used distances such as unweighted and weighted UniFrac [36].

Let *T* be the ORB-tree associated with a phylogenetic tree (e.g. the 97*%* Greengenes tree), and *S* the covariance matrix of *T*. In the context of phylogenetic informed metrics, environments are represented as probability mass functions over the OTUs (i.e. leaves). We denote those functions with lower-case letters such as *a* and *b*, and interpret them as probability models over *L*. In particular, *a* : *L* → [0, *+*∞) satisfies that *∑* _*x*∈*L*_ *a*(*x*) = 1 and, for each *e* ∈ *E, a*(*e*) = *∑* _*x*∈*e*_ *a*(*x*). With this convention, the DPCoA distance between two environments *a* and *b* is defined as [45, 20]:

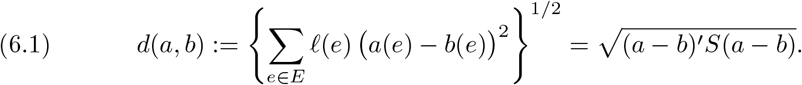

In particular, since *S* is positive definite, DPCoA corresponds to a Mahalanobis distance [38]; implying that *d*(·, ·) is a metric*—*in the mathematical sense*—*in ℝ^|*L*|^.

The weighted and unweighted UniFrac distances are instead defined as follows [36]:

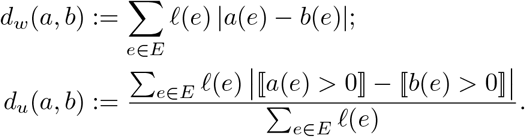

Both versions of UniFrac are known to satisfy the triangular inequality [37]. DPCoA is also more robust to unbiased noise but more sensitive to outliers than UniFrac [20]. Regardless of the metric of choice, the standard approach to linking environmental factors with microbial composition goes roughly as follows [34]. First, environmental samples are collected, and each environment is represented by its OTU composition on the leaves of the phylogeny of reference. Then, the pairwise distance matrix between the environments is computed, and the environments are embedded into a low-dimensional Euclidean space using standard techniques such as multidimensional scaling (MDS) [5]. Despite the noisy and high-dimensional nature of microbial datasets [53, 35, 24], this approach has been remarkably reliable for the ordination [31] of microbial environments in as little as 1-2 dimensions, and for correlating environmental factors with microorganisms. However, this approach does not usually explain correlations, which need to be justified by other means.

In what remains of this section, we apply our methods to the Greengenes phylogeny. First, we demonstrate significant sparsification of the associated covariance matrix after applying the Haar-like wavelet transform. Then, we motivate a new wavelet-based phylogenetic *β*-diversity metric corresponding to a multiscale analysis of the phylogenetic tree. Finally, we show that this wavelet-based metric can give novel insights into the relationship between environmental factors and OTU composition.

### 6.1. Greengenes Phylogenetic Covariance Matrix Sparsification

The 97*%* Greengenes tree has about 100,000 leaves. We can think of it as an ORB-tree by adding an external root and connecting it to the original root ° with a branch of length 0. (Alternatively, we could think of the Greengenes tree as two ORB-trees with their roots merged.) We denote the resulting ORB-tree as *T*.

The identity in equation (1.3) implies that the covariance matrix *S* of *T* is a 2 × 2 block diagonal matrix, with each block corresponding to an ORB-subtree. Approximately 94*%* of the almost 10 billion entries in *S* are non-zero because one of the ORB-subtrees (corresponding to the Archaea domain) is much smaller than the other*—*see Figure 6(a). This makes storing the covariance matrix of *T* challenging. Further, basic computational tasks such as finding the spectrum and inverting *S* for parameter estimation in phylogenetic comparative methods [29, 19] is infeasible because this large matrix is almost fully dense. We may use, however, the Haar-like matrix Φ associated with *T* to sparsify *S*. From Theorem 3.4, we can guarantee that *ζ* ≥ 0.9989, i.e. at least 99.89*%* of the entries in the similar matrix Φ′*S*Φ vanish. This significant compression of the matrix *S* can be appreciated in Figure 6(b).

**Fig. 6:**
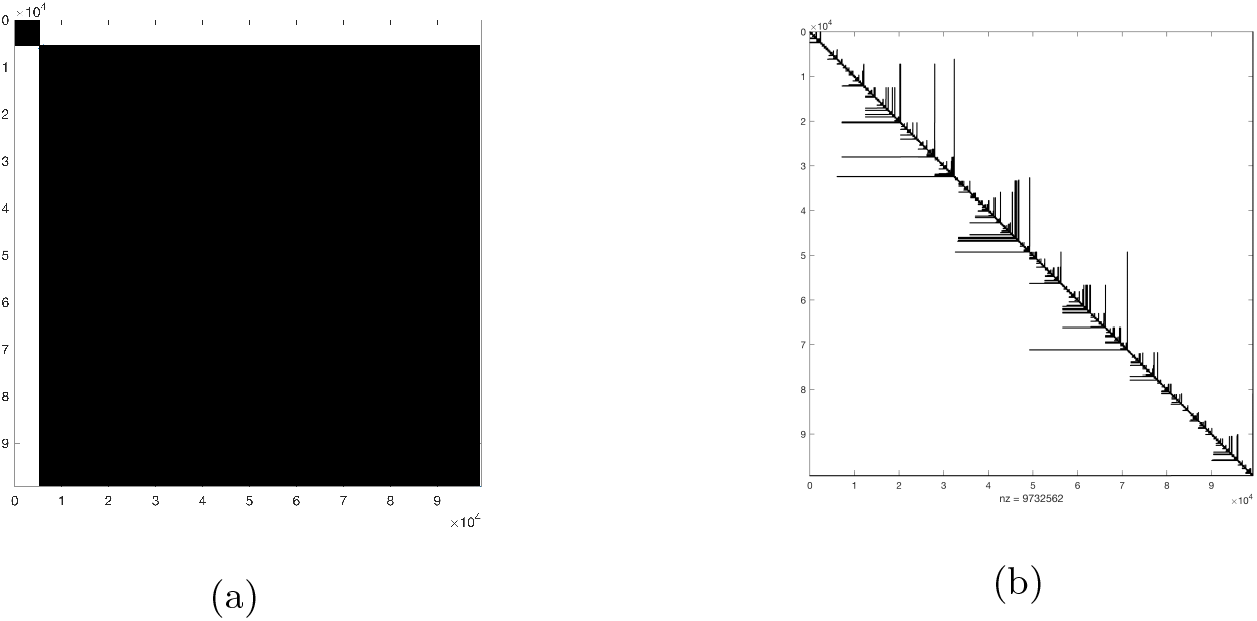
Heatmaps of matrices associated with the 97*%* Greengenes. Black (white) pixels denote non-zero (vanishing) entries. (a) Phylogenetic covariance matrix *S* of the 97*%* Greengenes tree. *S* has dimensions ∼ 10^5^× 10^5^. (b) Sparsified matrix Φ^*′*^*S*Φ.

We implemented Algorithm 2.1 using the sparse matrix packages from SciPy [58] to compute Φ^*′*^*S*Φ. As proof-of-principle, we used this compressed representation to compute the largest 500 eigenvalues of *S* to machine precision using SciPy’s implementation of the Lanczos algorithm. As seen in Figure 7, the eigenvalues of *S* decay rapidly. In fact, we found that *λ*_1_ (*S*) ∼ 1.27 × 10^5^, *λ*2(*S*) ∼ 4.75 × 10^3^, and trace(*S*) ∼ 1.65 × 10^5^, so the top-two eigenvalues already account for approximately 80*%* of the trace of *S*.

**Fig. 7:**
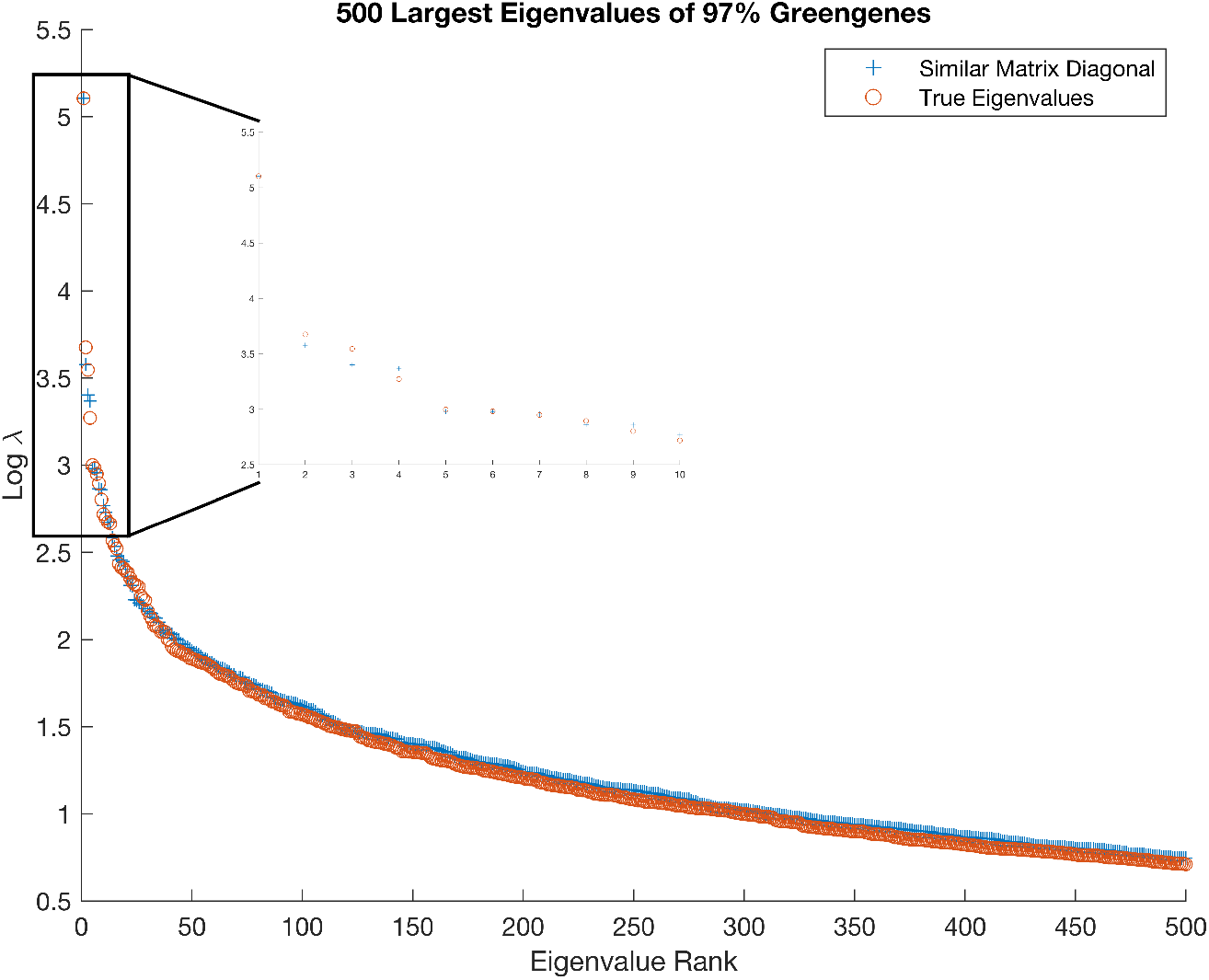
Spectrum decay of 97*%* Greengenes tree covariance matrix and corresponding approximation using Haar-like wavelets. Only the 500 most dominant eigenvalues of *S* are plotted as a function of their rank. Logarithms are in base-10.

As seen in Figure 7 also, the sorted diagonal entries in Φ^*′*^ *S*Φ (i.e. the quantities *λ* _*v*_, with *ν* ∈ *I*) approximate with ample accuracy the spectrum of *S*. For instance, max_*v*∈*I*_ *λ* _*v*_ underestimates *λ*1(*S*) with only about a 0.06*%* relative error. Anticipating this overall accuracy from *T* alone remains an open problem as neither our mathematical results, particularly Theorem 5.2, nor more general ones such as the Gershgorin’s circle theorem, Sylvester’s determinant theorem, and bounds found in [56, 7, 27] have been able to explain it.

### 6.2 A Wavelet Based Phylogenetic *β*-diversity Metric

Let *T* be the ORB-tree associated with a phylogenetic tree. Recall that *φv*, with *ν* ∈ *I*, is supported on *L*(*ν*), and together these functions form an orthonormal base of ℝ^|*L*|^. In particular, just as wavelets are traditionally used to localize signals at different scales, we may use the Haar-like basis of *T* to localize environmental OTU distributions on subsets of leaves defined by splits in the tree. This is particularly appealing from a biological standpoint. Indeed, the opposite signs of *φ* _*v*_ on the leaves of the left and right subtrees dangling from *ν* may be interpreted as a speciation event that conferred more fitness to present-day microorganisms descending from one of the subtrees than the other. We propose the following definition to convey these features into a phylogenetic *β*-diversity metric.

Recall that *λ* _*ν*_ = (Φ^*′*^*S*Φ)(*ν, ν*) *>* 0, for each *ν* ∈ *I*. Further, for a given environment *a* (i.e., OTU distribution over *L*), Φ^*′*^*a* is the projection of *a* onto the Haar-like basis of the reference tree.

#### Dfinition 6.1.

*The Haar-like distance between two environments a and b is the quantity*

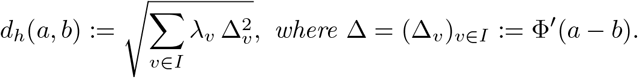

The specifics of this distance can be motivated as follows. On one hand, the terms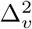, with *ν* ∈ *I*, convey the idea that *dh* regards two environments similar (different) when their OTU compositions project similarly (differently) onto the Haar-like basis of the reference tree. On the other hand, the weights *λ* _*v*_, with *ν* ∈ *I*, are motivated by the success of DPCoA in various biological investigations. To explain this, consider the matrices *D* := diag(*λ* _*v*_ : *ν* ∈ *I*) and *E* := Φ^*′*^*S*Φ − *D*. Observe that 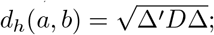 in particular, *dh* is a metric in ℝ^|*L*|^ because *D* is positive definite, and 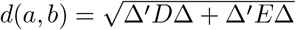. In large phylogenetic trees, however, we expect *E* to be mostly filled with zeroes due to Corollary 3.8 *—*which suggests considering *dh* as an alternative metric to DPCoA.

We have mentioned before that while traditional phylogenetic metrics (in conjunction with embedding techniques) have been remarkably successful at correlating microbial composition with environmental factors, these correlations cannot usually be explained from the metrics alone. The wavelet nature of the Haar-like distance has, however, the potential to explain said correlations. Indeed, the biological interpretation of the Haar-like basis conveyed by their sign flip suggests that if 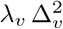 is comparatively large (small) for some *ν* ∈ *I*, then the speciation event associated with *ν* has a significant (little) influence differentiating the OTU distributions between two environments *a* and *b*. (There may be discrepancies between taxonomy and splits in a tree. In particular, while the aforementioned correlations may be explained by a phylogeny, they are not necessarily explained by a taxonomic classification.)

Previously we mentioned the equivalence of the Haar-like basis and ILR basis. Accordingly, compositional metrics with similar interpretations [42, 52] can be defined by projecting log-ratios of OTU counts onto the Haar-like basis (equivalently ILR basis). However, contrary to the Haar-like distance, these metrics do not account for phylogenetic induced covariance between OTUs.

### 6.3 Haar-like Distances of the Guerrero Negro microbial mat

A microbial mat is a bio-film of layered groups of microorganisms with coupled biochemistries. Their rich biodiversity, combined with the environmental gradients of light, oxygen, etc., offer an ideal setting to test phylogenetic *β*-diversity metrics.

The Guerrero Negro mat is hypersaline. It is located in Baja California Sur, Mexico. To demonstrate the insights possibly gained from the Haar-like distance, we applied it to a 16S rRNA data set of 18 samples at different depths of the Guerrero Negro mat [33, 22]. We used the 97*%* Greengenes as the reference phylogeny.

Earlier work [33] based on unweighted UniFrac showed a gradient of microbial composition in the mat with respect to depth*—*see top plot in Figure 8. (For a discussion regarding the “horseshoe” shape in the plot see [33, 43, 14].) As seen on the bottom plot of the same figure, we can practically reproduce this gradient using the Haar-like distance instead. Furthermore, as seen on the bottom two plots, the DPCoA and Haar-like distance produce nearly indistinguishable embeddings.

**Fig. 8:**
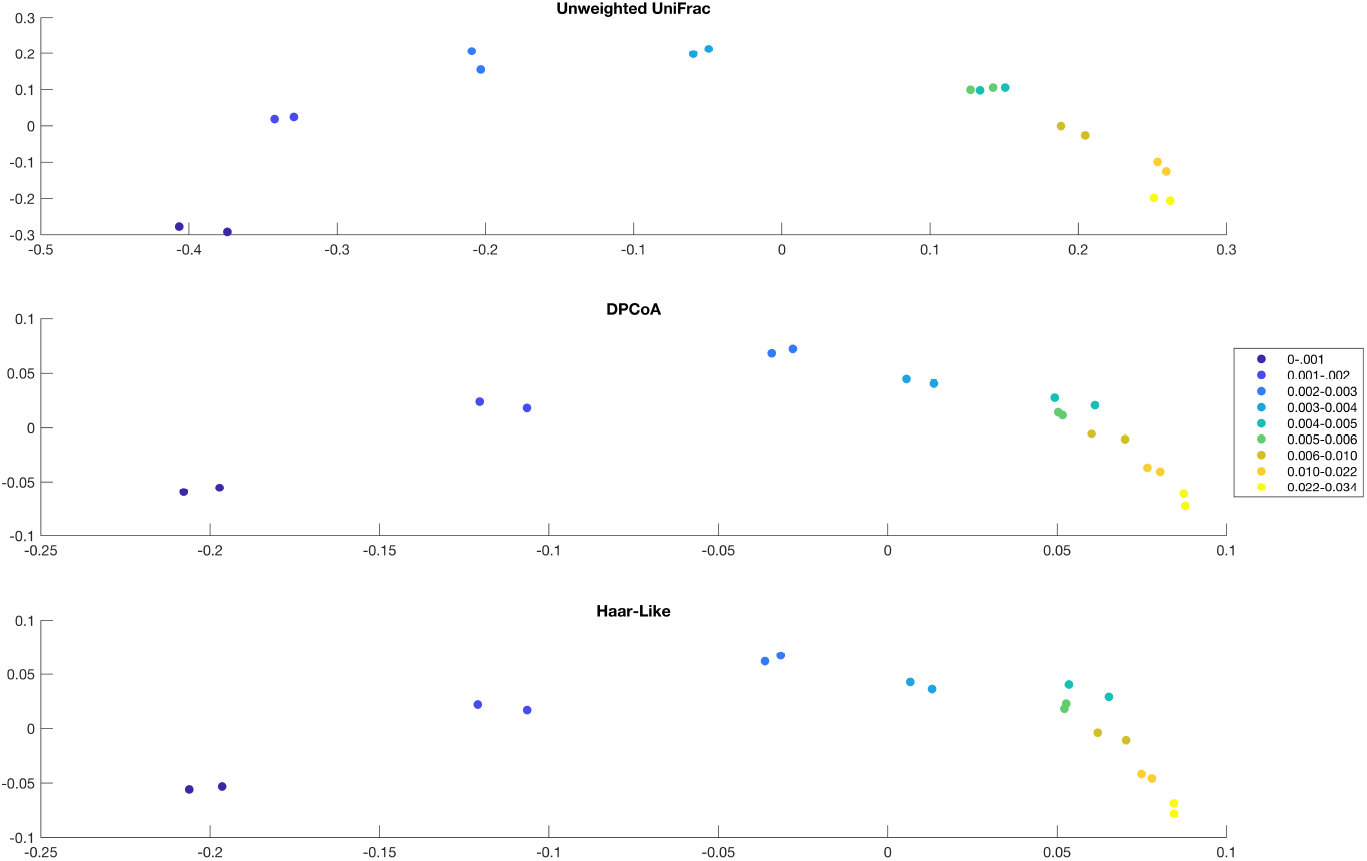
2-D MDS embeddings of samples from Guerrero Negro w.r.t. different metrics. The embeddings are based on unweighted UniFrac (top), DPCoA (middle), and the Haar-like distance (bottom). Depth varies from 0-0.034 meters.

While the three phylogenetic *β*-diversity metrics imply that mat depth drives a measurable change in OTU composition, we can go a step further with the Haar-like distance and determine which splits in the 97*%* Greengenes tree are responsible for this trend and quantify their importance. We demonstrate this by comparing the two extremes in the dataset: let *a* and *b* be the OTU compositions of the shallowest and deepest environment, respectively. Define Δ = Φ^*′*^(*b a*). Following the logic described in Section 6.2, we computed 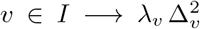, indexing interior nodes according to a postorder traversal of the 97*%* Greengenes tree. These values are shown in Figure 9. For our analysis, we focus on the largest 3 values, all of which are statistically significant (via the number of standard deviations they deviate from the mean). These are associated with the Haar-like wavelets 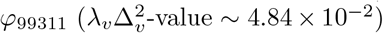, 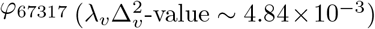, and 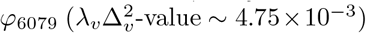. These correspond to splits at depths 10, 34, and 18 of the 97*%* Greengenes tree, respectively.

**Fig. 9:**
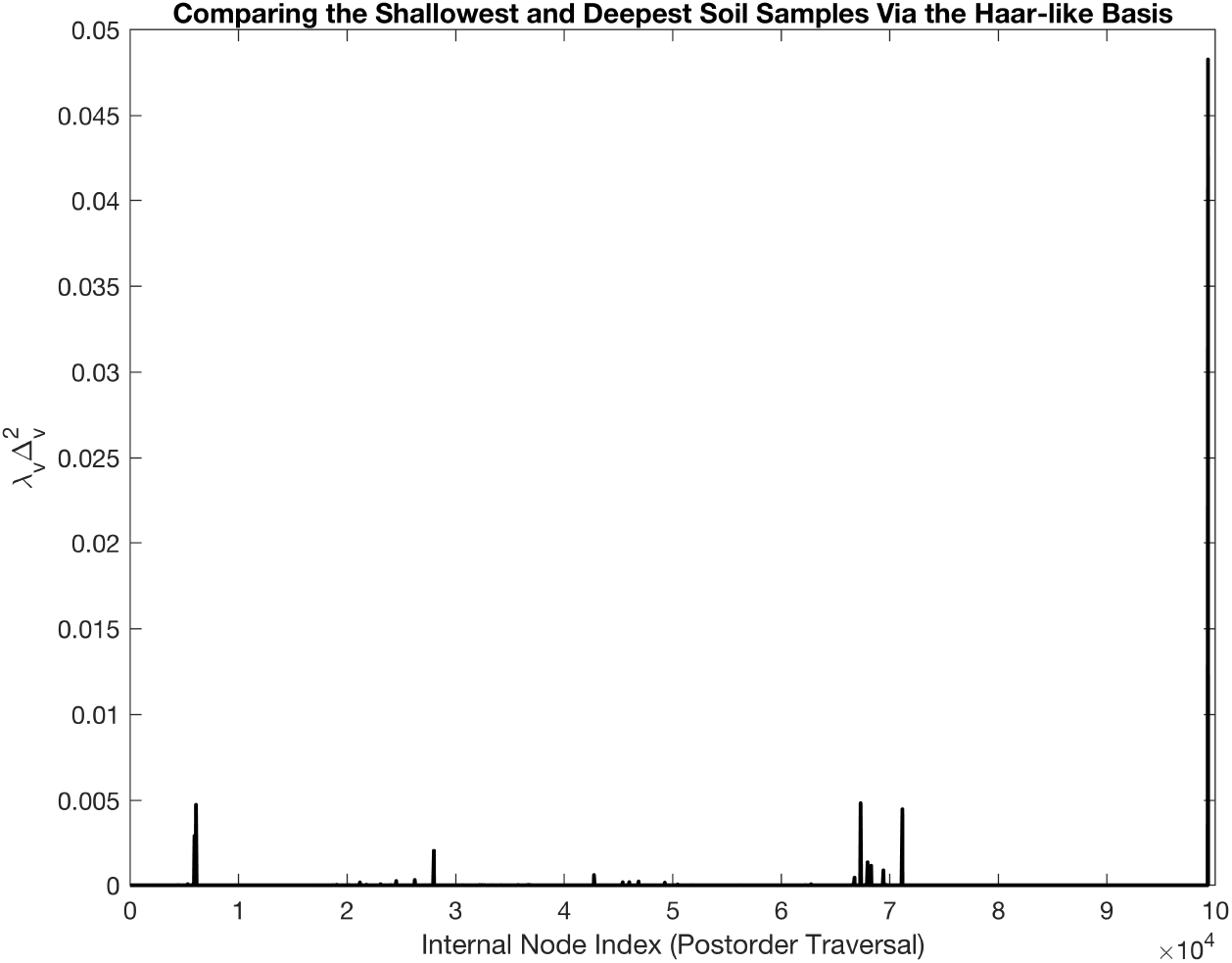
Plot of 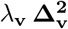, with *ν* ∈ *I*, to measure the Haar-like distance between the shallowest and deepest sample in the Guerrero Negro dataset. The average non-zero value of 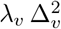 is ∼ 2.09 × 10^−5^. The standard deviation of these values is ∼ 7.55 × 10^−4^.

Notably, the split associated with *φ*_99311_ corresponds to the largest 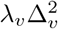-value. According to the associated taxonomic classification, the (say) left descendants of this split correspond to the phylum level classification of Cyanobacteria. This is consistent with the conclusion in [33], which correlated Cyanobacteria abundance changes with mat depth and explained the correlation by their ability to photosynthesize.

The other two wavelets provide novel insight into other important OTU composition differences driving the observed mat depth gradient in the Guerrero Negro dataset. Indeed, while the descendants of the split associated with *φ* _67317_ do not exhaust a taxonomic classification, all leaves under that split are classified as Anaerolineae. This differentiation between the shallowest and deepest sample may be due to Anaerolineae’s role as a anaerobic digester [59].

Finally, the split associated with *φ*_6079_ subdivides the Cyanobacteria phylum into further classes, including Oscillatoriophycideae, which is the third most abundant class of the Guerrero Negro dataset. The relevance of this split to differentiate shallow from deep samples may be explained by Oscillatoriophycideae’s photoautotrophic capability [55].

Our analysis of the Guerrero Negro mat shows that the Haar-like distance may be a valid alternative to other more common phylogenetic *β*-diversity metrics, primarily because it provides a systematic method for detecting statistically significant speciation events (and corresponding levels of OTU classification) that can link OTU composition with environmental factor gradients.

## 7. Conclusions

We have presented an approach to analyze and manipulate strictly ultrametric matrices through their ORB-tree representation. We demonstrated that the Haar-like wavelets associated with an ORB-tree provide an orthonormal basis with respect to which their associated matrix can be sparsified. Sparse representations may allow otherwise computationally infeasible but standard manipulations of these matrices, such as inverting and factoring them and characterizing their spectrum and eigenvectors. We also detailed a sparsification algorithm and showed that, with overwhelmingly high probability, only an asymptotically negligible fraction of the off-diagonal entries in random but large, strictly ultrametric matrices would remain non-zero after its application. Additionally, we provided some exact and approximate spectral results for ultrametric matrices based on the trace branch length “balancedness” of their ORB-tree. We also characterized the possible spectrums of strictly ultrametric matrices, giving further insight into the symmetric nonnegative inverse eigenvalue problem.

Conversely, the strictly ultrametric matrix associated with an ORB-tree corresponds to the phylogenetic covariance matrix of the tree. We applied our methods to the microbiologist’s Tree of Life covariance matrix as proof of concept. This covariance model is a standard reference in metagenomic studies, which rely on metrics such as UniFrac and Double Principal Coordinate Analysis (DPCoA). Motivated by the fact that the Tree of Life’s covariance matrix is significantly sparsified by the Haar-like wavelets, and that the diagonal of the sparsified matrix approximates with striking accuracy its spectrum, we introduced the Haar-like distance. Like the established metrics, this new metric measures the distance between pairs of microbial environments taking into account the relative abundance of microbes and their evolutionary relatedness. Unlike the established metrics, however, this new distance may be used to identify statistically significant speciation events linking microbial composition with environmental factors.

## Appendix A. Orthonormality of Haar-like bases

The statement that the Haar-like basis {*φ*_*v v*∈*I*_ associated with an ORB-tree is orthonormal is based on the concept of multiresolution analysis of Euclidean spaces in [21]. Here we justify this fact by first principles.

Let *u, ν* ∈ *I*. If *u* = *ν* then

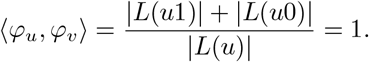

Instead, there are two possibilities when *u ≠ ν*. If *L*(*u*) ∩ *L*(*ν*) = ∅ then ⟨ *φv, φu*⟩ = 0 because *φ v* and *φu* have disjoint supports. Otherwise, if *L*(*u*) ∩ *L*(*ν*) ≠ ∅ then Lemma 2.4 let us assume without any loss of generality that *u* is an ancestor of *ν*. In particular, *L*(*ν*) ⊂ *L*(*u*) but also *φu* remains constant over *L*(*ν*). Therefore, for any given *x* ∈ *L*(*ν*):

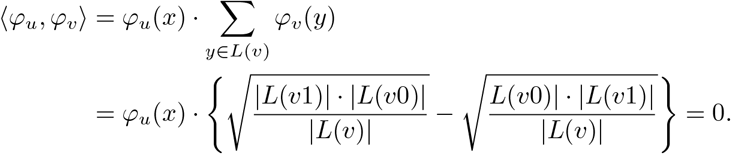

## Acknowledgments

This work has been partially funded by the NSF grant No. 1836914.

## Notes

https://github.com/edgor17/Sparsify-Ultrametric

### Competing Interest Statement

The authors have declared no competing interest.

### Summary of Updates

The manuscript has been revised after incorporating feedback from peers, people that contacted us after the first draft appeared in the bioRxiv and ArXiv, and an Associate Editor.

